# Characteristics of vaginal microbes and classification of the vaginal microbiome

**DOI:** 10.1101/2023.08.16.553525

**Authors:** Bin Zhu, Katherine M. Spaine, Laahirie Edupuganti, Andrey Matveyev, Myrna G. Serrano, Gregory A. Buck

## Abstract

**Background:** The vaginal microbiome (VMB) has been classified into several discrete community state types, some of which have been associated with adverse human health conditions. However, the roles of the many vaginal bacteria in modulating the VMB and health remain unclear.

**Methods:** The associations among the vaginal taxa and other vaginal taxa, the vaginal pH, and the host gene expression responses were determined by calculating the correlation among the relative abundance of the vaginal taxa, the association between the vaginal pH and the predominant taxon in the VMB, and the correlation between the relative abundance of the vaginal taxa and human gene expression at the transcriptional level, respectively. Using these associations, an alternative more informative method, the biological vagitype (BVT), is proposed to classify community state types of the VMB.

**Findings:** Most *Lactobacillus* spp., with the exception of *Lactobacillus iners*, show significant correlations with host gene expression profiles and negative associations with dysbiosis-associated vaginal taxa. Many non-*Lactobacillus* spp. exhibit varied correlations with *Lactobacillus* spp., the vaginal pH, and host gene expression. Compared to other dysbiotic taxa, including *Candidatus* Lachnocurva vaginae, *Gardnerella vaginalis* has a stronger positive correlation with vaginal pH and a stronger negative correlation with *Lactobacillus* spp. Most dysbiosis-associated taxa are associated with stress responses of the host at the transcriptional level, but the genus *Mycoplasma* has a uniquely strong positive correlation with host immune responses. The association between BVTs of the VMBs and host characteristics, e.g., race/ethnicity, microbial infection, smoking, antibiotics, high blood pressure, economic state, diet, and others, was examined. The BVT classification method improved overall performance in associating specific vaginal microbial populations with host characteristics and phenotypes.

**Interpretation:** This study sheds light on the biological characteristics of the vaginal microbiota, including some less abundant or still unculturable taxa. Since the BVT method was established based on these biological characteristics, the classification outcome of the VMB may have more clinical relevance. Because the BVT method performs better in associating specific vaginal community types with diseases, e.g., bacterial vaginosis and gonorrhea, it could be beneficial for the predictive modeling of adverse health.

**Funding:** This work was supported by grants [UH3AI083263, U54HD080784, and R01HD092415] from the National Institutes of Health; and support from the [GAPPS BMGF PPB] grant from the Global Alliance to Prevent Prematurity and Stillbirth. We would also like to thank the Office of Research on Women’s Health at NIH for their generous support.

**Research in context:** *Evidence before this study:* The vaginal microbiome (VMB) refers to the community of microorganisms in the female lower reproductive tract. The VMB is often a simple ecosystem dominated by a single species. The most predominant bacteria in the VMB include several *Lactobacillus* species and two non-*Lactobacillus* species, i.e., *Candidatus* Lachnocurva vaginae and *Gardnerella vaginalis. Lactobacillus* species produce lactic acid to lower the vaginal pH and inhibit the growth of disease-associated bacteria. Thus, the predominance of protective Lactobacilli, i.e., *L. crispatus, L. jensenii*, and *L. gasseri*, in the VMB is associated with overall vaginal health. However, the role of *L. iners* in promoting a healthy vaginal ecosystem is less clear. Actually, the biological and health relevance of many bacteria in the female lower reproductive tract is largely unknown. Some bacteria have low relative abundances, e.g., *Peptostreptococcus* and *Coriobacteriaceae* spp.; and others are not yet culturable, e.g., *Candidatus* Lachnocurva vaginae and BVAB TM7. When abundance of a taxon is low, its association with a host characteristic is a challenge. Previous methods to classify the VMB were based simply on their microbial compositions, and the biological characteristics of the vaginal bacteria were largely ignored. Thus, classification of these VMBs into biologically relevant community types, as described herein, should be helpful in determining their relevance to women’s reproductive health.

*Added value of this study:* This study examines three biological characteristics of bacteria in the VMB, i.e., the associations among different bacterial taxa, the vaginal pH, and the host response. Based on these three characteristics, the influence of these bacteria, particularly low abundant and unculturable bacteria, on vaginal health is evaluated. *L. iners* seems to be neutral in maintaining overall vaginal health. *Gardnerella vaginalis* is apparently more easily inhibited by *Lactobacillus* spp. than *Candidatus* Lachnocurva vaginae because of its stronger positive correlation with vaginal pH and negative correlation with *Lactobacillus*. The genus of *Mycoplasma* has a unique positive correlation with local immune responses, implying a role for *Mycoplasma* in promoting inflammation. Compared with previous methods to classify the VMB, a new method, considering the above three biological characteristics of bacteria in the VMB, has been established. The new method performs better in associating specific vaginal communities with host characteristics and phenotypes; e.g., bacterial vaginosis and gonorrhea.

*Implications of all the available evidence:* Accurate biological classification of the VMB is fundamental for assessing its impact on women’s health. Our classification scheme represents a step further toward that correct classification, eventually leading to new strategies for clinical assessment of the potential use of the VMB to diagnose or predict women’s reproductive health.

## Introduction

The vaginal microbiome (VMB) is generally a less complex ecosystem than the microbiomes in other human body sites, e.g., the oral cavity and the gastrointestinal tract^1^. Many VMBs are dominated by a single taxon^2–4^, and the predominance of several *Lactobacillus* spp., e.g., *L. crispatus, L. gasseri*, and *L. jensenii*, is a hallmark of overall vaginal health^5–7^. VMBs predominantly of these taxa are considered ‘optimal’^7,8^. VMBs dominated by non-*Lactobacillus* spp., e.g., *Gardnerella vaginalis, Candidatus* Lachnocurva vaginae (bacterial vaginosis-associated bacterium 1 or BVAB1), and other anaerobes, or a mixture of multiple anaerobes, are considered dysbiotic and nonoptimal. These latter VMBs are often associated with adverse gynecological and obstetric outcomes. VMBs dominated by *L. iners* are considered transitional or intermediate with relevance to women’s gynecologic and obstetric health^8^.

Production of lactic acid is an essential mechanism by which protective lactobacilli, e.g., *L. crispatus, L. gasseri*, and *L. jensenii*, maintain the health of the female lower reproductive tract. Lactic acid decreases the vaginal pH, inhibits the growth of dysbiosis-associated microbes^9,10^, and interacts with the human immune system to diminish pro-inflammatory responses that could be induced by dysbiosis-associated microbes^11,12^. Thus, a lower vaginal pH is associated with a higher relative abundance of these *Lactobacillus* spp. in the VMB^3,13^ and lower risks of microbial infections in the female reproductive tract and adverse outcomes in pregnancy, e.g., bacterial vaginosis (BV)^14^ and preterm birth^15^. In contrast to VMBs dominated by the protective lactobacilli, VMBs dominated by *L. iners* are associated with a lower concentration of D-Lactic acid in the vaginal fluid^16^ and a higher vaginal pH^3^. Furthermore, *L. iners* produces the virulence factor inerolysin which can disrupt the integrity of the epithelial barrier in vitro^17^, has a stronger potential than *L. crispatus* for stimulating host pro-inflammatory responses in vitro^18^, and is associated with increased risks of preterm birth and miscarriage^19,20^. Hence, the contribution of *L. iners* to overall vaginal health is less positive and more transitional than that of the more protective *Lactobacillus* spp^21^.

The human immune system can be stimulated by dysbiosis-associated bacterial taxa. Levels of several pro-inflammatory cytokines and chemokines, e.g., IL-1_β_, IL-6, IL-8, RANTES, G-CSF, and TNF-_α_, are increased by *G. vaginalis* and *Atopobium vaginae* in human epithelial cell models^18,22^. Dysbiosis-associated taxa also generate virulence factors, e.g., mucinase, sialidase, vaginolysin, glycosulfatase, and others, that can disrupt the mucosal and epithelial barriers^17,23–25^. Thus, it is not surprising that the dominance of dysbiosis-associated taxa in the VMB increases the risks of BV, sexually transmitted infections (STIs), adverse outcomes in pregnancy, and other adverse conditions of women’s reproductive health^23,25–28^.

Biofilm formation of *G. vaginalis* is enhanced in co-cultures with *A. vaginae, Fusobacterium nucleatum*, and other common vaginal taxa^29^, and ammonia produced by *Prevotella bivia* promotes the growth of *G. vaginalis*^30^. Hence, interactions among these dysbiosis-associated taxa may increase the competitiveness of these taxa in the female lower reproductive tract.

Efforts have been made to associate specific vaginal taxa with women’s adverse health conditions. However, at least partially because of limited sample sizes, sequencing depth, and participants’ differences in age, genetic background, economic state, and other host factors, not to mention consistent descriptions of the various adverse health conditions, the identification of specific disease-associated taxa often fails to reach significance. However, the classification of the VMB into distinct groups has permitted the association of these groups with adverse conditions of health^5^ as well as age^31^, race^2^, pregnancy^32^, menstrual stage^33^, and others.

Hierarchical clustering of dissimilarity among 16S rRNA taxonomic profiles has revealed five common community state types (CSTs I-V) dominated by *L. crispatus, L. gasseri, L. iners*, one or multiple anaerobic taxa, and *L. jensenii*, respectively^3^. CST IV can be further classified into several subtypes^4^. CST IV has been associated with increased risks of BV^34^, STIs^35^, preterm birth^20,32,36^, and other adverse health conditions. An alternative approach categorizes VMBs into vagitypes based on the most abundant taxon in a VMB^2,28^. Previous works have illustrated that protective *Lactobacillus*-dominated vagitypes and community state types are enriched in pregnancy^2,7,28,32^ and transition of the VMB to *Ca*. Lachnocurva vaginae vagitype during pregnancy is associated with a higher risk of preterm birth^28^. However, the biological differences among taxa in the VMB are generally not considered in these classification methods. In clinical settings, a few methods, e.g., Amsel’s criteria and the Nugent Score^37^, are used to associate the VMB with adverse conditions and risk of adverse outcomes by quantifying taxa with specific morphotypes, measuring the vaginal pH and assessing host responses, e.g., abnormal vaginal discharge and odor. These methods are semi-quantitative, somewhat subjective, and generally lack species-level resolution.

Herein, we explore differences among bacteria populations in the lower female reproductive tract, the association of taxa in the VMB with the vaginal pH and host responses, and the association among taxa in the VMB. These associations, coupled with vagitype classifications, were used to develop a more informative strategy for the classification of the VMB.

## Methods

### Cohort

Samples used in this study were from two large cohorts. The first included 2881 sets of high-quality 16S rRNA gene sequencing data targeting the V1-V3 variable regions and related demographic and clinical metadata (SI Data 2 Sheet 2 row 6) collected from vaginal samples of 2881 reproductive-age women under the umbrella of the Vaginal Human Microbiome Project (VaHMP)^38–40^. The second cohort included 130 sets of 16S rRNA sequence data targeting the same V1-V3 variable region as above and the human gene expression profiles generated from vaginal samples of 130 pregnant women collected in the first visit in the Multi-Omic Microbiome Study-Pregnancy Initiative (MOMS-PI) project^28,38^ (SI Data 1 Sheet 1).

### Data processing

After quality control, trimming, merging paired sequence reads, and removing human reads as previously described^28^, high-quality sequences of the 16S rRNA amplicons collected in the MOMS-PI and VaHMP projects were aligned to the STIRRUPS database^41^ using ublast^42^ as previously described^2,28^ for taxonomic assignment of vaginal taxa at species level. Because many vaginal taxa at the species level had low relative abundances in the vaginal microbiome, low abundant taxa were merged into taxa at the genus level as previously reported^4^. Only taxa with a relative abundance of no less than 1% on average in the vaginal microbiomes in the VaHMP dataset were kept at the species level. For example, four *Lactobacillus* spp., i.e., *L. crispatus, L. iners, L. gasseri*, and *L. jensenii*, with relative abundance no less than 1%, were kept in the analyses. All other *Lactobacillus* taxa were merged as ‘Other *Lactobacillus*’. None of *Streptococcus* spp. had a relative abundance higher than 1%, and those taxa were combined as ‘*Streptococcus*’. Taxa with at least 0.1% (or 0.01%) relative abundance in at least 5% (or 15%) of the samples were kept in the feature table of the 16S rRNA profiles as previously described^2,28^. Samples in the feature table with total sample reads less than 5,000 were excluded from this study, leaving 130 and 2881 samples from the MOMS-PI and VaHMP projects, respectively. Paired metatranscriptomic data of the host cells in the MOMS-PI project and metadata in the VaHMP project were also involved in this study. The high-quality metatranscriptomic sequences generated in the MOMS-PI project were aligned to the transcriptome of Homo sapiens (assembly GRCh38.p14) downloaded from the NCBI GenBank database using Bowtie 2^43^ for producing a feature table of the host gene expression.

### Classification of the vaginal microbiome by community state types

The feature table of the 16S rRNA profiles was converted to a relative abundance table. Samples in the relative abundance table were clustered by calculating the Euclidean distance and using the ‘complete’ method as previously described^3^ in the ‘pheatmap’ function in R. The VMBs were classified into nine CSTs using the ‘cutree’ function in R.

### Classification of the vaginal microbiome by vagitypes

The VMBs were assigned to vagitypes according to the most abundant species with a relative abundance of 30% or greater, as previously described^2,28^. A VMB with the relative abundance of the most abundant taxon less than 30% was assigned to ‘No type’.

### Diversity analysis

The feature table of the 16S rRNA profiles was normalized by rarefaction to the depth of the lowest number of reads in a sample (>5,000). Alpha diversity was quantified by calculating the Shannon index using the ‘vegan’ package in R^44^. The difference in the alpha diversity was measured by the Wilcoxon test. Beta diversity was evaluated by Bray-Curtis distance using the ‘vegan’ package in R. The median values of the Bray-Curtis distances between the VMBs with two different vagitypes were clustered by calculating the Euclidean distance and using the ‘ward.D’ method in the ‘pheatmap’ function in R. The difference in the beta diversity between two vagiypes was tested by the ‘adonis2’ function, a PERMANOVA analysis, in the ‘vegan’ package in R. An alternative method to investigate the beta diversity of the VMBs was to normalize the feature table of the 16S rRNA profiles by the centered log-ratio transformation. The beta diversity was quantified and visualized by a t-SNE of Bray-Curtis distances using the ‘Rtsne’ package in R^45^.

### Association among taxa in the vaginal microbiome

The feature table of the 16S rRNA profiles was normalized by the centered log-ratio transformation. The association between each taxa dyad was measured by Spearman’s correlation. The coefficient and significance of a correlation were quantified by an R- and *P*-value, respectively. The *P*-values were adjusted by the Benjamini-Hochberg procedure using the ‘adjust.p’ function in the ‘cp4p’ package in R. The R-values with matched adjusted *P*-values larger than 0.05 were replaced by zeros to remove insignificant correlations in the following analysis. Taxa in the VMB were clustered according to the R-values of the Spearman’s correlation using the ‘pheatmap’ function with default settings in R and were classified using the ‘cutree’ function in R.

### The impact of the vaginal pH on the vaginal microbiome

There were 2037 VMBs collected in the VaHMP project with values of the vaginal pH in the matched metadata. These VMBs were divided into pH high, medium, and low groups, with sample numbers evenly distributed in the three groups. Beta diversity of the VMBs was quantified and visualized by a t-SNE of Bray-Curtis distances using the ‘Rtsne’ package in R, and the difference of the beta diversity associated with the vaginal pH was measured by the ‘adonis2’ function in the ‘vegan’ package in R. Differential abundance analysis was performed using the ‘ADLEx2’ package^46^ in R. The adjusted *P*-value of relative abundance differences was tested by the ‘aldex.ttest’ function using the Mann–Whitney U test value, followed by the Benjamini-Hochberg correction. The relative abundance change was measured by the ‘aldex.effect’ function and quantified by the per-feature median difference between two conditions.

### The association between the vaginal taxa and the vaginal pH

The feature table of the 16S rRNA profiles was normalized by the centered log-ratio transformation, and the association between the vaginal taxa and the vaginal pH was tested by linear regression using the ‘lm’ function in R. The *P*-values of the linear correlations were adjusted by the Benjamini-Hochberg procedure. Alternatively, the median value of the vaginal pH was calculated when a taxon was dominated in the VMBs with a relative abundance of no less than 30%.

### Host responses to taxa in the vaginal microbiome

The feature table of the 16S rRNA profiles was normalized by the centered log-ratio transformation, and the feature table of the host gene expression was normalized by a variance stabilizing transformation using the ‘varianceStabilizingTransformation’ function in the ‘DESeq2’ package^47^ in R. The association between the vaginal taxa and the host genes was measured by Spearman’s correlation. The R- and *P*-values were adjusted as described in ‘Association among taxa in the vaginal microbiome’ in the Methods. The taxa and host genes were both clustered by calculating the Euclidean distances and using the ‘ward.D’ method in the ‘pheatmap’ function in R. The vaginal taxa and host genes were classified into nine and five groups, respectively, using the ‘cutree’ function in R.

### Functional enrichment of the host gene clusters

Genes belonging to each gene cluster were input into the DAVID database^48^. Three DAVID categories, i.e., UP_KW_BIOLOGICAL_PROCESS, GOTERM_BP_DIRECT, and KEGG_PATHWAY, were chosen for the functional enrichment analysis. *P*-values of the functional enrichment were adjusted by the Benjamini-Hochberg procedure.

### Classification of the vaginal microbiomes by biological vagitypes

The vaginal taxa were classified based on three characteristics, i.e., correlation with other taxa in the VMB, the vaginal pH when a taxon was dominated in the VMB, and correlation with the host gene expression. The clustering of the vaginal taxa according to correlations with other taxa and the host gene expression are categorical values and the median values of the vaginal pH when a taxon is dominated in the VMB are numeric values. These three types of values were combined into a matrix. Dissimilarity among the vaginal taxa was evaluated by calculating the Gower distance using the ‘daisy’ function with the ‘metric’ parameter equal to ‘gower’ in the ‘cluster’ package in R. The vaginal taxa were classified into seven groups using the Gower distances and the “complete” clustering method in R. Taxa that were predominant in less than four VMBs were not involved in this analysis. The VMBs were assigned to BVT I to VII according to the vagitypes and the classification of the vaginal taxa. The VMBs without classification were assigned to ‘BVT others’.

### Host characteristics

There are three types of values in the metadata (SI Data 2). The first type of host characteristic contains binary values -‘Yes’ and ‘No’, e.g., ‘caucasian’, ‘high_blood_pressure_status’, and ‘bv’. The second type has ordinal values containing more than two but less than seven levels, e.g., ‘income’, ‘education’, and ‘alcohol_12months’. These values were converted to two levels, ‘High’ and ‘Low’, with similar sample numbers in the two groups. More details are in the custom codes ‘VaHMP.R’ at lines 1505-1634. The third type of host characteristic has numeric values with more than or equal to seven levels, e.g., ‘sample_pH’, ‘agesex’, and ‘BMI’.

### Comparison of the vaginal microbiome classification using the CST, vagitype, and BVT methods

The VMBs were assigned to CSTs, vagitypes, and BVTs, respectively. In SI Data 2 sheet 2, the proportions of participants with a particular host characteristic (for metadata with binary values) or a high level of a host characteristic (for metadata with ordinal values) in each specific VMB type classification or the median (25% quantile, 75% quantile) values of a particular host characteristic in each specific VMB type classification (for metadata with numeric values) are shown. For example, ‘37.9%; 488’ in column D row 8 in SI Data 2 sheet 2 represents that, in the participants whose information about ‘African American’ have been collected, there are 488 participants with VMBs of CST I type and 37.9% of the 488 participants are African American. ‘63.6%; 464’ in column G row 8 represents that, in the participants whose information about ‘income’ have been collected, there are 464 participants with VMBs of CST I type and 63.6% of the 464 participants are at a high level of income. ‘4.4 (4, 4.7); 389’ in column B row 8 represents that, in the participants whose information about ‘sample_ph’ have been collected, there are 389 participants with VMBs of CST I type and the median (25% quantile, 75% quantile) values of the sample pH for the 389 participants are 4.4 (4, 4.7). For Pearson’s Chi-squared tests as described below, the host characteristics with numeric values were also converted to two levels, ‘High’ and ‘Low’, with similar sample numbers in the two groups. The difference among the distribution of each host characteristic in different VMB types was examined by Pearson’s Chi-squared tests with *P*-values adjusted by the Benjamini-Hochberg procedure. Thus, a lower *P*-value indicates that the matched host characteristic is more distinguished among different types of VMBs. For example, ‘7.05E-83’ in column B row 17 in SI Data 2 sheet 2 represents that, in the participants whose information about ‘sample_ph’ have been collected, the vaginal pH levels are significantly different among the CST categories. The performances of the CST, vagitype, and BVT methods on classifying the VMBs were evaluated by pairwisely comparing the adjusted *P*-values of the Pearson’s Chi-squared tests using the Mann–Whitney U test.

### Ethics

Participants were enrolled in the Multi-Omic Microbiome Study: Pregnancy Initiative (MOMS-PI) and the Vaginal Human Microbiome Project under the umbrella of the National Institutes of Health Human Microbiome Project (https://commonfund.nih.gov/hmp). Women were enrolled in women’s clinics associated with the Virginia Commonwealth University Health Center. Study protocols were approved by the Virginia Commonwealth University institutional review board under protocols IRB# HM12169 or HM15527. Written informed consent or parental permission and assent were provided by participants or minors older than 15 years, respectively. Exclusion criteria included women incapable of understanding the informed consent or assent forms or who were incarcerated. Demographic, health histories, dietary assessments, and clinical data (e.g., gestational age, height, weight, blood pressure, vaginal pH, diagnosis, etc.) were collected. Other exclusion criteria included: 1) inability to self-sample due to any reason; 2) significant vaginal bleeding; 3) ruptured membranes; 4) herpes lesions.

## Results

### The vaginal microbiome classified by CSTs and vagitypes

The 16S rRNA taxonomic profiles of 2881 women collected in the Vaginal Human Microbiome Project (VaHMP)^38–40^ were classified into CSTs by hierarchical clustering and as vagitypes by measuring predominant taxa (Fig. 1a). Because most VMBs consist of a single predominant taxon, these two methods produced similar results, particularly in the VMBs with CST I (*L. crispatus* vagitype), CST II (*L. gasseri* vagitype), CST III (*L. iners* vagitype), CST V (*L. jensenii* vagitype), and CST IVB (*G. vaginalis* vagitype). CST IVA is mainly composed of VMBs with *Ca*. Lachnocurva vaginae, *A. vaginae*, and *Mycoplasma* vagitypes. CST IVC is complex in composition and overlaps with multiple vagitypes, e.g., *Sneathia amnii, Streptococcus* spp., *P. bivia*, No Type (see Methods), and other vagitypes.

**Fig. 1.**
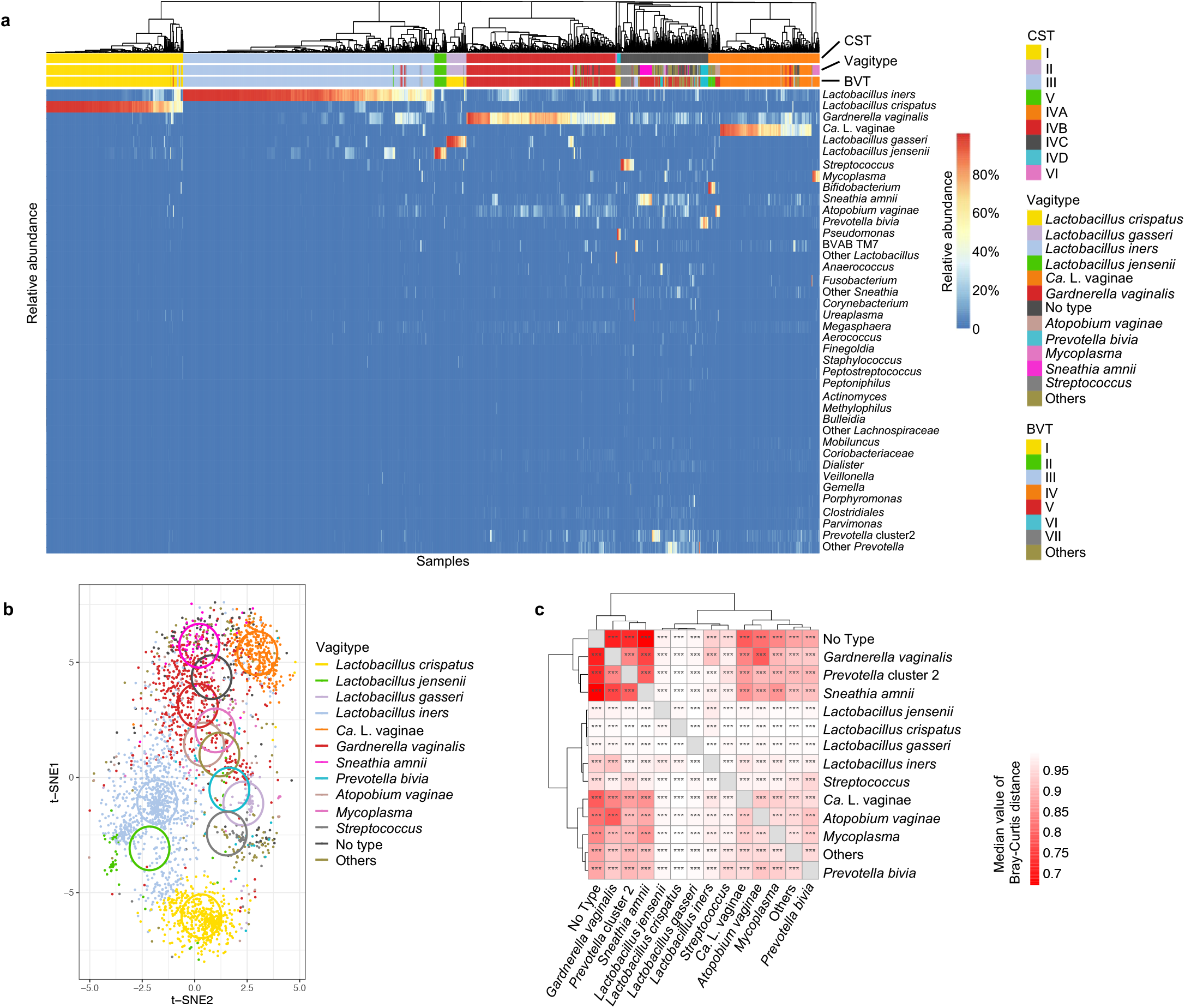
Classification and dissimilarity of the vaginal microbiome. **(a)** The heatmap illustrates the composition of the 2881 vaginal microbiomes (VMBs) collected in the VaHMP project. The VMBs were classified by the community state type (CST) method and were color-coded according to the classifications of the VMBs by three methods (see Methods), i.e., CST, vagitype, and biological vagitype (BVT). **(b)** The dissimilarity of the VMBs was quantified and visualized by a t-distributed stochastic neighbor embedding plot (t-SNE). The VMBs were color-coded by the vagitypes. The circle indicates the center of the VMBs with a specific vagitype on the plot. **(c)** A heatmap shows the dissimilarity of the VMBs evaluated and clustered according to the median values of the Bray-Curtis distances between each vagitype dyad. Additionally, pairwise analysis of the difference among the vagitypes was performed using the Adonis test. The *P*-values of the Adonis test indicated by asterisks are shown on the heatmap. *** *P*-value ≤ 0.001. Ca. L. vaginae represents *Candidatus* Lachnocurva vaginae.

Pairwise analysis of the alpha diversities associated with each vagitype showed that most exhibited significant differences (Fig. S1a). Consistent with previous observations^49^, all VMBs with *Lactobacillus* vagitypes were less complex than those with non-*Lactobacillus* vagitypes. Dissimilarity of the 2881 VMBs classified by vagitypes was visualized by a t-distributed stochastic neighbor embedding (t-SNE) plot (Fig. 1b and SI Movie 1) and a heatmap showing the clustering of these vagitypes based on median values of Bray-Curtis distance between each vagitype dyad (Fig. 1c). Both the t-SNE plot and the pairwise Adonis tests offered on the heatmap indicate that each of these vagitypes is significantly different in beta diversity from each of the other vagitypes, illustrating the rationale of the vagitype method (Fig. 1b and c). The within-group Bray-Curtis distance of VMBs with *L. crispatus* vagitype is lower than that of other *Lactobacillus* vagitypes (Fig. S1b) and the relative abundance of *L. crispatus* in VMBs with *L. crispatus* vagitype is higher than other taxa in their matched vagitypes (Fig. S1c), suggesting that *L. crispatus* tends to exclude other taxa and implying a stronger impact of *L. crispatus* on modulating the VMBs.

Significant differences exist in the VMB classifications according to the CST and vagitype strategies. For example, the *Streptococcus* vagitype is similar to the *Lactobacillus* vagitypes on the t-SNE plot (Fig. 1b) and the heatmap (Fig. 1c) but is closer to non-*Lactobacillus* vagitypes in the hierarchical clustering (Fig. 1a). Similarly, the *A. vaginae* vagitype clusters with the *Ca*. Lachnocurva vaginae vagitype in the hierarchical clustering and heatmap but is closer to the *G. vaginalis* vagitype on the t-SNE plot. These inconsistencies suggest a more in-depth study of the biological characteristics of the vaginal taxa to improve and more functionally characterize the VMBs.

### Associations among bacteria in the vaginal microbiome

Associations among bacterial taxa in the VMBs were measured by calculating Spearman’s correlation of their relative abundances. Hierarchical clustering of these correlations separated the vaginal taxa into six main groups (Fig. 2a). Group A includes most *Lactobacillus* spp. (*L. crispatus, L. gasseri, L. jensenii*, and other *Lactobacillus* spp. (see Methods)) as well as less common taxa including *Corynebacterium, Bifidobacterium, Pseudomonas, Veillonella, Staphylococcus*, and *Methylophilus. L. iners*, included in Group B with *Ureaplasma, Streptococcus*, and some other taxa, is the only *Lactobacillus* species not included in Group A.

**Fig. 2.**
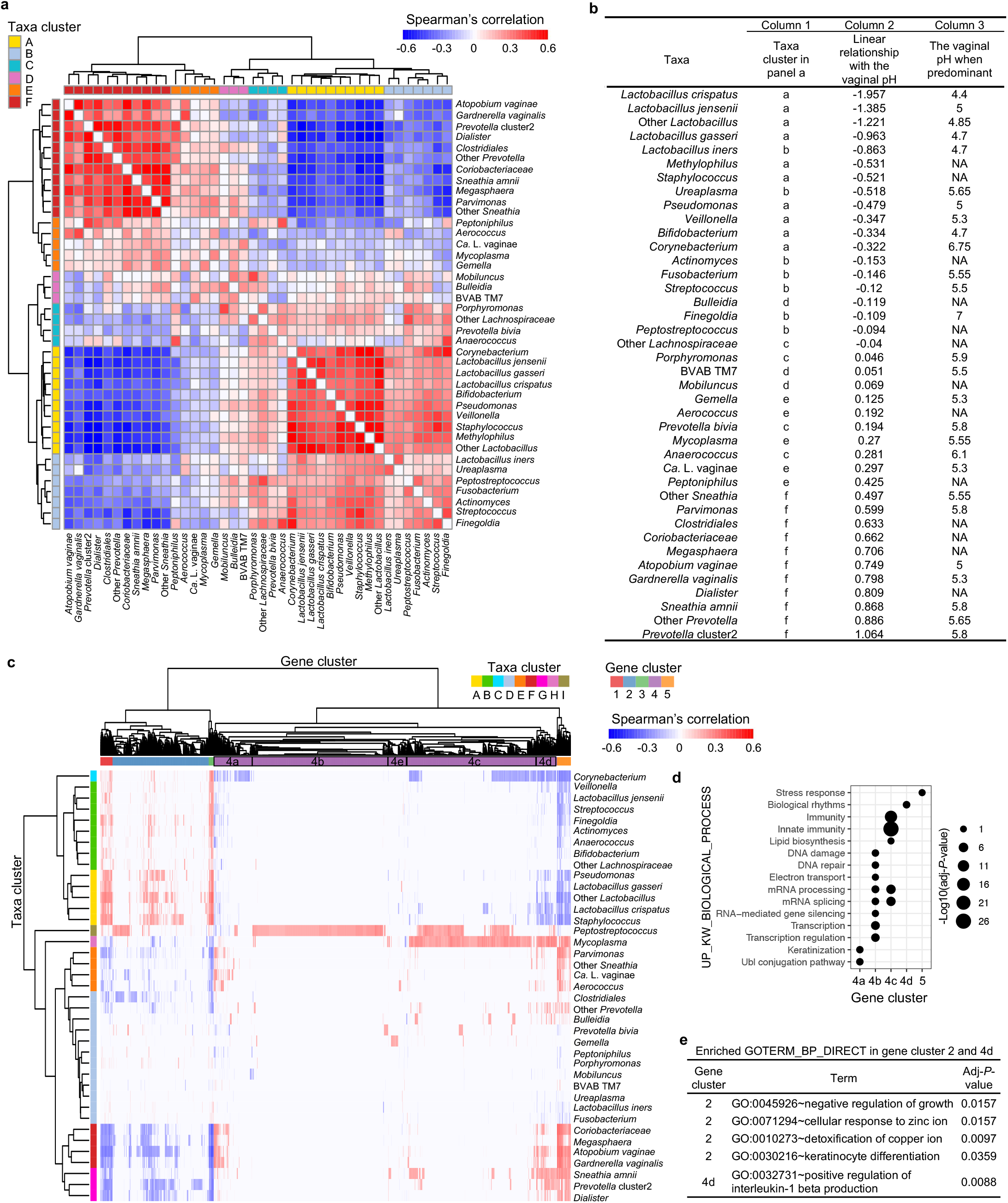
Characterization of taxa in the vaginal microbiome. **(a)** The association among the vaginal taxa was quantified by Spearman’s correlation between the relative abundance of each taxa dyad. The correlation coefficients with adjusted *P*-values larger than 0.05 were replaced by zeros. The clustering of the correlation coefficients is shown in the heatmap, and the classification of the vaginal taxa according to the clustering results is color-coded. **(b)** The slopes of the linear regressions between the relative abundance of a vaginal taxon and the vaginal pH and the median vaginal pH, when a taxon is predominant, are shown (see details in Fig. S2 and Table S1). **(c)** The association between the relative abundance of the vaginal taxa and the levels of individual host genes was quantified by Spearman’s correlation. The correlation coefficients with adjusted *P*-values larger than 0.05 were replaced by zeros. The clustering of the correlation coefficients is shown by a heatmap, and the classifications of the vaginal taxa and the host genes according to the clustering results are color-coded. **(d)** Functional enrichment of genes in each host gene cluster analyzed by the DAVID database with functions annotated by the ‘UP_KW_BIOLOGICAL_PROCESS’ category is shown. **(e)** Functional enrichment of genes in the host gene clusters 2 and 4d analyzed by the DAVID database with functions annotated by the GOTERM_BP_DIRECT category is shown (see more pathway enrichment results in SI Data 1 Sheet 4).

Overall, taxa in Group A were positively correlated with taxa in Groups B, C, and D but negatively correlated with taxa in Groups E and F (Fig. 2a). The overall negative association between Groups A and F was more robust than that between Groups A and E, implying a more potent inhibition of taxa in Group F by *Lactobacillus* spp. Consistent with an apparent lower exclusionary effect^16–20^, *L. iners* had weaker negative correlations with dysbiosis-associated taxa than other *Lactobacillus* spp. (Fig. 2a). *A. vaginae* had the strongest positive correlation with *G. vaginalis*, consistent with the reported synergistic effect of *A. vaginae* and *G. vaginalis* in biofilm formation^29^.

### Association between taxa in the vaginal microbiome and the vaginal pH

Since a low vaginal pH is generally associated with women’s reproductive health^7^, the association between the relative abundance of taxa in the VMB and the vaginal pH was evaluated by linear regression and the degree of the linear relationship was quantified by the slope of the linear regression (Fig. 2b and S2). Taxa that were less abundant at lower vaginal pH generally also exhibited a stronger negative correlation with protective Lactobacilli, implying that these taxa could be more greatly inhibited by lactic acid (Fig. 2b Column 2 versus 1). Some non-*Lactobacillus* spp. found in Group A in the association study above (Fig. 2a), e.g., *Bifidobacterium, Pseudomonas, Fusobacterium, Staphylococcus, Streptococcus*, and *Ureaplasma*, also showed a higher relative abundance with a decrease of the vaginal pH (Fig. 2b). Conversely, taxa found in Groups C, D, and E, which generally included taxa associated with dysbiosis and adverse health indications, largely exhibited an increased relative abundance in samples exhibiting a high pH. Similar results were observed in the differential abundance analysis among VMBs with different levels of vaginal pH using ALDEx2^46^ (Fig. S3). It is worth noting that several dysbiosis-associated taxa, e.g., *G. vaginalis* and *A. vaginae*, only show significant abundance differences between the VMBs with low and medium levels of vaginal pH but not between those with medium and high levels of vaginal pH (Fig. S3c). This result implies that *G. vaginalis* and *A. vaginae* are more sensitive to the vaginal pH at a range lower than 4.5. However, several taxa that are weakly associated with the vaginal pH in the linear regression analysis, e.g., *Ca*. Lachnocurva vaginae and BVAB TM7, are also not significantly associated with the vaginal pH in the differential abundance analysis (Fig. S3c).

Because several vaginal bacteria, e.g., *Ca*. Lachnocurva vaginae and BVAB TM7, are still not readily culturable, the ability of each taxon to lower the vaginal pH was estimated by testing the vaginal pH when the taxon was predominant in the VMB. The results showed that the predominance of *L. crispatus* in the VMB was most tightly related to a low vaginal pH (Fig. 2b). Several non-*Lactobacillus* taxa, e.g., *Streptococcus, Ureaplasma*, and *Corynebacterium* spp., showed a negative linear correlation with the vaginal pH but were similar to *Ca*. Lachnocurva vaginae and *G. vaginalis* in the ability to lower the vaginal pH. This pattern is consistent with the hypothesis that the sensitivity of a vaginal taxon to the vaginal pH rather than its ability to reduce the vaginal pH is driving its association with other vaginal taxa in the VMB (compare column 1 to 2 and 3 in Fig. 2b).

### Host responses to bacteria in the vaginal microbiome

Host transcriptional levels in the lower female reproductive tract in samples exhibiting different VMB profiles were measured by Spearman’s correlation between the relative abundance of taxa in the VMB and host gene expression in the sample. Hierarchical clustering of the correlations identified nine bacterial taxa groups and five human gene groups (Fig. 2c and SI Data 1). Human gene cluster 4 was partitioned into subgroups 4a-4e. Most *Lactobacillus* spp., except for *L. iners*, were classified in taxon clusters A or B and generally correlated with opposite host responses compared to taxa in clusters E-H containing dysbiosis-associated taxa. *L. iners* was grouped in taxon cluster D with *Ureaplasma*, BVAB TM7, *Mobiluncus, Fusobacterium, P. bivia*, and others that exhibited limited correlations with the host gene expression. *Mycoplasma* and *Peptostreptococcus*; comprising taxon clusters H and I, respectively, were associated with specific host expression profiles compared to other dysbiosis-associated taxa.

Functional enrichment of the human gene clusters was tested by the DAVID analysis tool^48^, and the results of three DAVID categories, i.e., UP_KW_BIOLOGICAL_PROCESS, GOTERM_BP_DIRECT^50^, and KEGG_PATHWAY^51^, are listed (SI Data 1). The expression of gene clusters 4a, 4d, and 5 with functions enriched in the ubiquitin-like protein (Ubl) conjugation pathway, positive regulation of IL-1_β_ production, and stress responses, respectively, was positively correlated with taxon clusters E-G, containing dysbiosis-associated taxa including *Ca*. Lachnocurva vaginae, *Sneathia* spp., *G. vaginalis, Prevotella* cluster 2, and *Dialister* spp. In contrast, expression of these gene clusters was negatively correlated with protective *Lactobacillus, Corynebacterium*, and several non-Lactobacillus taxa (Fig. 2c-e) that are positively correlated with *Lactobacillus* (see above, Fig. 2a). The Ubl conjugation pathway is involved in the autophagy of host cells in the inflammatory response^52^. Gene cluster 4b, associated with RNA modulation and DNA damage and repair, was mainly positively correlated with *Peptostreptococcus*, and *Peptostreptococcus* and *Mycoplasma* species were positively correlated with Gene cluster 4c, which includes genes involved in innate immunity.

Keratinocyte differentiation was influenced by both protective *Lactobacillus* spp. (taxon clusters A and B, Fig. 2c and e) and dysbiosis-associated taxa; e.g., *G. vaginalis, A. vaginae*, and *Ca*. Lachnocurva vaginae (taxon clusters E and F, Fig. 2c and d). However, most genes correlated with protective *Lactobacillus*; e.g., genes encoding corneodesmosin (CDSN)^53^, desmoplakin (DSP)^54^, and cystatin A (CSTA)^55^, are different from those correlated with dysbiosis-associated taxa; e.g., genes encoding small proline-rich (SPRR) proteins, involucrin (IVL), and late cornified envelope 3E (LCE3E) (SI Data 1 Sheet 4 column G). The latter set of genes is mainly associated with the production of keratin and SPRR proteins that can protect the epidermal barrier from bacterial disruption^56,57^. Thus, dysbiosis-associated taxa and more protective taxa seem to have different influences on the keratinization of human cells at the transcriptional level.

### Classification of taxa in the vaginal microbiome by biological vagitypes

Three characteristics, i.e., the pattern of association among taxa in the VMB, the vaginal pH when the VMB is predominantly a single taxon, and host responses to a taxon at the transcriptional level, were applied to cluster twenty vaginal taxa into seven groups using Gower’s distance (Fig. 3a). *L. crispatus* clustered with *L. gasseri*, close to a cluster including *L. jensenii*, but more distantly related to a group including *L. iners. G. vaginalis* was grouped with *A. vaginae*, close to a cluster including *Sneathia* spp. and *Prevotella* cluster2. Additionally, *Ca*. Lachnocurva vaginae was in the same group as *Mycoplasma*, and *P. bivia* clustered with BVAB TM7.

**Fig. 3.**
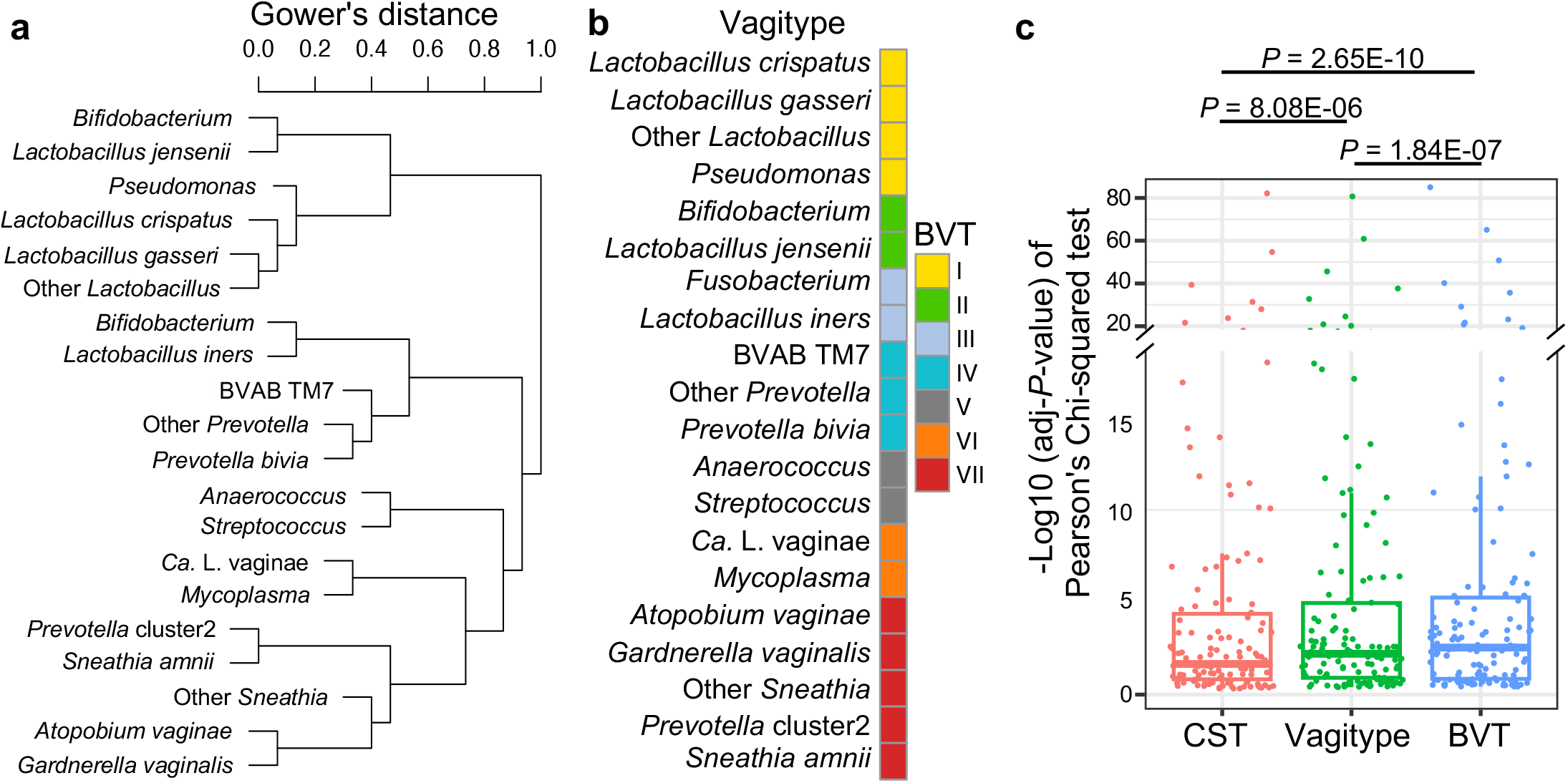
A new classification method for the vaginal microbiome. **(a)** Three types of values, i.e., the categories of the clustering of the vaginal taxa in the associations with other taxa in the VMB and with the host gene expression, and the median value of the vaginal pH when a taxon was dominated in the VMB, were applied to classify the vaginal taxa into seven groups by calculating the Gower’s distance. **(b)** The vagitypes with matched vaginal taxa clustered together are assigned to the same biological vagitype (BVT). Unclassified VMBs are assigned to ‘BVT others’. **(c)** The performance of the CST, vagitype, and BVT methods in associating specific community state types with host characteristics was quantified by Pearson’s Chi-squared test. The pairwise comparisons of the adjusted *P*-values of the Pearson’s Chi-squared test generated by the three methods using the Mann–Whitney U test are shown.

We have implemented the above classification scheme to classify vagitypes into seven biological vagitypes (BVT I to BVT VII) (Fig. 3b). An additional ‘BVT others’ includes all other unclassified VMBs. Most of these ‘BVT others’ are ‘No_type’ classified by vagitypes, representing complex VMBs lacking an obviously predominant taxon. The proportion of participants with a particular host characteristic in each specific VMB type classification was assessed, and the difference among the distribution of each host characteristic in different VMB types was determined by the Pearson’s Chi-squared tests with *P*-values adjusted by the Benjamini-Hochberg procedure (SI Data 2, see Methods for detailed explanation). By comparing the adjusted *P*-values, BVT seemed to perform better overall in associating specific community types of the VMB with the host characteristics than the original CST or vagitype strategies (Fig. 3c).

Notably, non-*Lactobacillus* spp. in BVTs I, II, and V were distinguished from those in other BVTs in host response (Fig. 2c). Consistent with this observation, BVTs I, II, and V were associated with a lower risk of BV (SI Data 2). Despite being associated with similar levels of vaginal pH (Fig. 2b), BVT V but not BVTs IV, VI, VII, and VIII was generally associated with a lower risk of BV, consistent with the hypothesis that activation of the host immune and other systems rather than high vaginal pH is critical in the pathogenesis of BV.

Women with VMBs of BVT VI, which includes *Ca*. Lachnocurva vaginae and *Mycoplasma* spp., have the highest proportion of BV, trichomoniasis, sexual activity, pregnancy, lower levels of income and height, the youngest age of the first sex, etc. (SI Data 2). Additionally, 85.1% of women with VMBs of BVT VI are African Americans. This result is consistent with a higher risk of BV in Black women^7^. Abnormal vaginal odor and discharge and vaginal douching were most frequently observed in the women with VMBs of BVT others, implying an association of these host characteristics with a high complexity of the VMB.

## Discussion

In this study, taxa with similar linear correlation with the vaginal pH were more likely clustered together in the bacterial association network (Fig. 2b), implying that the vaginal pH is an essential parameter that mediates associations among taxa in the VMB^7^. Likely due to the high resistance of some non-*Lactobacillus* species, e.g., *Bifidobacterium, Pseudomonas, Corynebacterium*, and *Staphylococcus*, to a low vaginal pH, these taxa clustered together with protective *Lactobacillus* spp. in the bacterial association network (Fig. 2a). Other factors, e.g., a synergistic effect of *G. vaginalis* and *A. vaginae* in biofilm formation^29^, could also impact the bacterial associations in the VMB.

In the association between the vaginal taxa and the host gene transcriptional levels, some well-known potentially pathogenic genera, e.g., *Ureaplasma, Pseudomonas, Streptococcus*, and *Staphylococcus*, exhibited no or even negative correlations with the expression of host stress or immune response-related genes, implying that the interaction between these bacteria and the host in the lower reproductive tract is different from that of taxa in BVT VI and VII. A plausible explanation for this observation is that these genera include different species or strains with disparate virulence factors. Alternatively, these taxa may have mechanisms to prevent, or at least avoid, inducing host defenses.

Since taxa of lower abundance were predominant in only very few samples, we were unable to accurately assess the vaginal pH when they were predominant (Table S1). Therefore, the BVT method only considers the twenty most abundant taxa in these samples (Fig. 3a). However, using our criteria, 95.6% of the 2881 VMBs studied herein were assigned to BVT I through BVT VII.

In contrast to the classification of the VMBs by CST or vagitype, the BVT strategy clusters the *L. crispatus* and *L. gasseri* VMBs together in BVT I. Similarly, the *G. vaginalis, A. vaginae*, and *S. amnii* dominant VMBs were also included in the same BVT VII (Fig. 3b). This result is consistent with the fact that taxa in these groups correlated positively with each other in relative abundance (Fig. 2a) and were associated with similar host responses (Fig. 2c). Consistent with the CST classification, both *Mycoplasma* and *Ca*. Lachnocurva vaginae-predominant VMBs are in BVT VI (Fig. 1a and 3b) mainly because of their similar pattern in the bacterial correlation network and a similar association with a lower vaginal pH (Fig. 2).

Compared to other *Lactobacillus* spp., *L. iners* showed a weaker negative correlation with dysbiosis-associated taxa (Fig. 2a) and had limited interaction with the host gene expression (Fig. 2c). Thus, it seems that *L. iners* is neutral relative to other *Lactobacillus* spp. in the maintenance of overall vaginal health according to results in this study. These observations support the hypothesis that VMBs dominated by *L. iners* are transitional or intermediate with relevance to women’s gynecologic and obstetric health^8^.

The association between the vaginal taxa and host responses was studied using metatranscriptomics data of the VMB from the MOMS-PI cohort^28,38^ which includes only pregnant women. Although the BVT method works well in the study (Fig. 3c), another metatranscriptomics dataset containing non-pregnant women could potentially improve the BVT method. The large sample size of the VaHMP cohort, including 2881 sets of high-quality VMBs and metadata, enhanced the statistical power of related analyses. However, a larger cohort with more information on the vaginal pH when a taxon was predominant could improve the BVT method.

The BVT strategy possibly provides a more clinically relevant stratification of VMBs. Thus, taxa with common biological functions are more likely classified together. Interestingly, the BVT method may begin to clarify the contribution of different taxa to adverse health conditions. Thus, bacterial vaginosis, for which classification remains controversial, seems to be more common in women with BVT VI microbiome profiles, whereas women with symptoms of discharge and odor are more likely to have the more complex microbiome under the umbrella of BVT others. Continued study is required to further clarify these relationships between VMBs and women’s health.

Since the BVT method is based on the vagitype method, the community type classified by BVT is a definite value, not depending on the clustering result in a dataset. Thus, the classification of a VMB by the BVT method is not impacted by other samples in the same dataset and could be beneficial for the predictive modeling of adverse health.

## Contributors

G. A. B., M. G. S., and B. Z. conceived and designed the study. B. Z. performed the analyses and wrote the paper. K. M. S. and A. M. participated in sample collection and sequencing. L. E. performed pretreatment of the raw sequencing data. M.G.S. and G.A.B. edited the paper.

## Acknowledgment

We gratefully acknowledge the participants in the VaHMP and MOMS-PI projects who generously provided samples to these studies. We also acknowledge the contributions of all members of the Vaginal Microbiome Consortium at VCU, whose participation made this report possible. We would also like to thank the Office of Research on Women’s Health, National Institute of Child Health Development, National Institute of Allergy and Infectious Disease, National Human Genome Research Institute, at the National Institutes of Health, and the Global Alliance to Prevent Prematurity and Stillbirth in Seattle for their generous support. The sequence data included in this study were generated in the Genomics Core at Virginia Commonwealth University. Analysis was performed on servers provided by the Center for High-Performance Computing at Virginia Commonwealth University. This work was supported by grants [UH3AI083263, U54HD080784, and R01HD092415] from the National Institutes of Health; and support from the [GAPPS BMGF PPB] grant from the Global Alliance to Prevent Prematurity and Stillbirth. We would also like to thank the Office of Research on Women’s Health at NIH for their generous support.

## Data sharing statement

Raw 16S rRNA sequences and controlled-access metadata of the VaHMP project^38–40^ are available from the Sequence Read Archive at NCBI (project ID phs000256). Metadata collected in the VaHMP project and applied in this study are shown in SI Data 2. Raw 16S rRNA sequences and limited metadata of the MOMS-PI project^28,38^ are available from the HMP DACC (https://portal.hmpdacc.org). Controlled-access data, including raw metatranscriptomic sequences and metadata of the MOMS-PI project, are available at National Center for Biotechnology Information’s controlled-access dbGaP (study no. 20280; accession ID phs001523.v1.p1) and the SRA under BioProject IDs PRJNA326441, PRJNA326442, and PRJNA326441. Further information and requests for resources should be directed to and will be fulfilled by the lead contact, Gregory A. Buck. All the codes are available on GitHub (https://github.com/GregoryBucklab/Vaginal_taxa_classification).

## References

1. Consortium, H. M. P. Structure, function and diversity of the healthy human microbiome. nature 486, 207 (2012).

2. Serrano, M. G. et al. Racioethnic diversity in the dynamics of the vaginal microbiome during pregnancy. Nat. Med. 25, 1001–1011 (2019).

3. Ravel, J. et al. Vaginal microbiome of reproductive-age women. Proc. Natl. Acad. Sci. 108, 4680–4687 (2011).

4. VALENCIA: a nearest centroid classification method for vaginal microbial communities based on composition. vol. 8 (2020).

5. Greenbaum, S., Greenbaum, G., Moran-Gilad, J. & Weintraub, A. Y. Ecological dynamics of the vaginal microbiome in relation to health and disease. Am. J. Obstet. Gynecol. 220, 324–335 (2019).

6. Anahtar, M. N., Gootenberg, D. B., Mitchell, C. M. & Kwon, D. S. Cervicovaginal Microbiota and Reproductive Health: The Virtue of Simplicity. Cell Host Microbe 23, 159–168 (2018).

7. Zhu, B., Tao, Z., Edupuganti, L., Serrano, M. G. & Buck, G. A. Roles of the Microbiota of the Female Reproductive Tract in Gynecological and Reproductive Health. Microbiol. Mol. Biol. Rev. MMBR e0018121 (2022) doi:10.1128/mmbr.00181-21.

8. Zheng, N., Guo, R., Wang, J., Zhou, W. & Ling, Z. Contribution of Lactobacillus iners to Vaginal Health and Diseases: A Systematic Review. Front. Cell. Infect. Microbiol. 11, 792787 (2021).

9. Aldunate, M. et al. Antimicrobial and immune modulatory effects of lactic acid and short chain fatty acids produced by vaginal microbiota associated with eubiosis and bacterial vaginosis. Front. Physiol. 6, 164 (2015).

10. Tyssen, D. et al. Anti-HIV-1 Activity of Lactic Acid in Human Cervicovaginal Fluid. mSphere 3, p(2018).

11. Hearps, A. C. et al. Vaginal lactic acid elicits an anti-inflammatory response from human cervicovaginal epithelial cells and inhibits production of pro-inflammatory mediators associated with HIV acquisition. Mucosal Immunol. 10, 1480–1490 (2017).

12. Delgado-Diaz, D. J. et al. Distinct immune responses elicited from cervicovaginal epithelial cells by lactic acid and short chain fatty acids associated with optimal and non-optimal vaginal microbiota. Front. Cell. Infect. Microbiol. 9, 446 (2019).

13. Miller, E. A., Beasley, D. E., Dunn, R. R. & Archie, E. A. Lactobacilli dominance and vaginal pH: why Is the human vaginal microbiome unique? Front. Microbiol. 7, 1936 (2016).

14. Ceccarani, C. et al. Diversity of vaginal microbiome and metabolome during genital infections. Sci. Rep. 9, 14095 (2019).

15. Hauth, J. C. et al. Early pregnancy threshold vaginal pH and Gram stain scores predictive of subsequent preterm birth in asymptomatic women. Am. J. Obstet. Gynecol. 188, 831–835 (2003).

16. Witkin, S. S. et al. Vaginal biomarkers that predict cervical length and dominant bacteria in the vaginal microbiomes of pregnant women. MBio 10, e02242–19 (2019).

17. Ragaliauskas, T. et al. Inerolysin and vaginolysin, the cytolysins implicated in vaginal dysbiosis, differently impair molecular integrity of phospholipid membranes. Sci. Rep. 9, 10606 (2019).

18. Anton, L. et al. Common cervicovaginal microbial supernatants alter cervical epithelial function: mechanisms by which Lactobacillus crispatus contributes to cervical health. Front. Microbiol. 9, 2181 (2018).

19. Kindinger, L. M. et al. The interaction between vaginal microbiota, cervical length, and vaginal progesterone treatment for preterm birth risk. Microbiome 5, 6 (2017).

20. Chang, D.-H. et al. Vaginal microbiota profiles of native Korean women and associations with high-risk pregnancy. J. Microbiol. Biotechnol. 30, 248–258 (2020).

21. Petrova, M. I., Reid, G., Vaneechoutte, M. & Lebeer, S. Lactobacillus iners: friend or foe? Trends Microbiol. 25, 182–191 (2017).

22. Doerflinger, S. Y., Throop, A. L. & Herbst-Kralovetz, M. M. Bacteria in the vaginal microbiome alter the innate immune response and barrier properties of the human vaginal epithelia in a species-specific manner. J. Infect. Dis. 209, 1989–1999 (2014).

23. Wiggins, R., Hicks, S. J., Soothill, P. W., Millar, M. R. & Corfield, A. P. Mucinases and sialidases: their role in the pathogenesis of sexually transmitted infections in the female genital tract. Sex. Transm. Infect. 77, 402–408 (2001).

24. Mohammadzadeh, R., Sadeghi Kalani, B., Kashanian, M., Oshaghi, M. & Amirmozafari, N. Prevalence of vaginolysin, sialidase and phospholipase genes in Gardnerella vaginalis isolates between bacterial vaginosis and healthy individuals. Med. J. Islam. Repub. Iran 33, 85 (2019).

25. Roberton, A. M. et al. A novel bacterial mucinase, glycosulfatase, is associated with bacterial vaginosis. J. Clin. Microbiol. 43, 5504–5508 (2005).

26. Ilhan, Z. E., Łaniewski, P., Tonachio, A. & Herbst-Kralovetz, M. M. Members of Prevotella genus distinctively modulate innate immune and barrier functions in a human three-dimensional endometrial epithelial cell model. J. Infect. Dis. 222, 2082–2092 (2020).

27. Gelber, S. E., Aguilar, J. L., Lewis, K. L. T. & Ratner, A. J. Functional and phylogenetic characterization of Vaginolysin, the human-specific cytolysin from Gardnerella vaginalis. J. Bacteriol. 190, 3896–3903 (2008).

28. Fettweis, J. M. et al. The vaginal microbiome and preterm birth. Nat. Med. 25, 1012–1021 (2019).

29. Castro, J., Machado, D. & Cerca, N. Unveiling the role of Gardnerella vaginalis in polymicrobial Bacterial Vaginosis biofilms: the impact of other vaginal pathogens living as neighbors. ISME J. 13, 1306–1317 (2019).

30. Pybus, V. & Onderdonk, A. B. Evidence for a commensal, symbiotic relationship between Gardnerella vaginalis and Prevotella bivia involving ammonia: potential significance for bacterial vaginosis. J. Infect. Dis. 175, 406–413 (1997).

31. Ouarabi, L. et al. Vaginal microbiota: age dynamic and ethnic particularities of Algerian women. Microb. Ecol. 82, 1020–1029 (2021).

32. DiGiulio, D. B. et al. Temporal and spatial variation of the human microbiota during pregnancy. Proc. Natl. Acad. Sci. 112, 11060–11065 (2015).

33. Langner, C. A. et al. The vaginal microbiome of nonhuman primates can be only transiently altered to become Lactobacillus dominant without reducing inflammation. Microbiol. Spectr. 9, e0107421 (2021).

34. Fudaba, M., Kamiya, T., Tachibana, D., Koyama, M. & Ohtani, N. Bioinformatics analysis of oral, vaginal, and rectal microbial profiles during pregnancy: a pilot study on the bacterial co-residence in pregnant wWomen. Microorganisms 9, p(2021).

35. Shannon, B. et al. Association of HPV infection and clearance with cervicovaginal immunology and the vaginal microbiota. Mucosal Immunol. 10, 1310–1319 (2017).

36. Tabatabaei, N. et al. Vaginal microbiome in early pregnancy and subsequent risk of spontaneous preterm birth: a case–control study. BJOG Int. J. Obstet. Gynaecol. 126, 349–358 (2019).

37. Nugent, R. P., Krohn, M. A. & Hillier, S. L. Reliability of diagnosing bacterial vaginosis is improved by a standardized method of gram stain interpretation. J. Clin. Microbiol. 29, 297–301 (1991).

38. Integrative HMP (iHMP) Research Network Consortium. The Integrative Human Microbiome Project: dynamic analysis of microbiome-host omics profiles during periods of human health and disease. Cell Host Microbe 16, 276–289 (2014).

39. The Integrative Human Microbiome Project. Nature 569, 641–648 (2019).

40. Fettweis, J. et al. The vaginal microbiome: disease, genetics and the environment. Nat. Preced. 1–1 (2010).

41. Fettweis, J. M. et al. Species-level classification of the vaginal microbiome. BMC Genomics 13 Suppl 8, S17 (2012).

42. Edgar, R. C. Search and clustering orders of magnitude faster than BLAST. Bioinforma. Oxf. Engl. 26, 2460–2461 (2010).

43. Langmead, B. & Salzberg, S. L. Fast gapped-read alignment with Bowtie 2. Nat. Methods 9, 357–359 (2012).

44. Jari Oksanen et al. vegan: community ecology package. (2020).

45. Van der Maaten, L. & Hinton, G. Visualizing data using t-SNE. J. Mach. Learn. Res. 9, (2008).

46. Fernandes, A. D., Macklaim, J. M., Linn, T. G., Reid, G. & Gloor, G. B. ANOVA-like differential expression (ALDEx) analysis for mixed population RNA-Seq. PloS One 8, e67019 (2013).

47. Love, M. I., Huber, W. & Anders, S. Moderated estimation of fold change and dispersion for RNA-seq data with DESeq2. Genome Biol. 15, 550 (2014).

48. Sherman, B. T. et al. DAVID: a web server for functional enrichment analysis and functional annotation of gene lists (2021 update). Nucleic Acids Res. 50, W216–221 (2022).

49. Mehta, S. D. et al. Host genetic factors associated with vaginal microbiome composition in Kenyan women. mSystems 5, p(2020).

50. Mi, H., Muruganujan, A., Ebert, D., Huang, X. & Thomas, P. D. PANTHER version 14: more genomes, a new PANTHER GO-slim and improvements in enrichment analysis tools. Nucleic Acids Res. 47, D419–D426 (2019).

51. Ogata, H. et al. KEGG: Kyoto Encyclopedia of Genes and Genomes. Nucleic Acids Res. 27, 29–34 (1999).

52. Cadwell, K. Crosstalk between autophagy and inflammatory signalling pathways: balancing defence and homeostasis. Nat. Rev. Immunol. 16, 661–675 (2016).

53. Dubash, A. D. & Green, K. J. Desmosomes. Curr. Biol. 21, R529–R531 (2011).

54. Boyden, L. M. et al. Dominant de novo DSP mutations cause erythrokeratodermia-cardiomyopathy syndrome. Hum. Mol. Genet. 25, 348–357 (2016).

55. Blaydon, D. C. et al. Mutations in CSTA, encoding Cystatin A, underlie exfoliative ichthyosis and reveal a role for this protease inhibitor in cell-cell adhesion. Am. J. Hum. Genet. 89, 564–571 (2011).

56. Geisler, F. & Leube, R. E. Epithelial Intermediate Filaments: Guardians against Microbial Infection? Cells 5, p(2016).

57. Zhang, C. et al. Small proline-rich proteins (SPRRs) are epidermally produced antimicrobial proteins that defend the cutaneous barrier by direct bacterial membrane disruption. eLife 11, p(2022).

