## Supplementary materials for "Characteristics of vaginal microbes and classification of the vaginal microbiome"

a

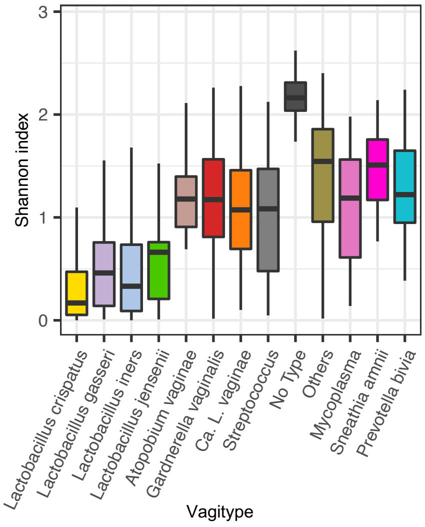


| | Lactobacillus iners | Lactobacillus gasseri | Others | Streptococcus | Gardnerella vaginalis | Ca. L. vaginae | Atopobium vaginae | Lactobacillus jensenii | Sneathia amnii | No Type | Prevotella bivia | Mycoplasma |
|---|---|---|---|---|---|---|---|---|---|---|---|---|
| Lactobacillus crispatus | 1.26E-10 | 1.94E-05 | 1.86E-48 | 6.43E-15 | 1.20E-120 | 4.10E-81 | 1.88E-17 | 2.91E-06 | 1.84E-29 | 4.30E-62 | 2.11E-17 | 2.15E-15 |
| Lactobacillus iners | | 2.70E-01 | 1.35E-41 | 1.10E-09 | 7.37E-111 | 5.60E-63 | 3.87E-13 | 3.01E-02 | 8.83E-26 | 1.29E-67 | 1.55E-13 | 6.87E-11 |
| Lactobacillus gasseri | | | 6.76E-19 | 4.25E-07 | 3.02E-26 | 3.06E-19 | 1.63E-11 | 2.29E-01 | 1.61E-19 | 2.32E-32 | 1.06E-11 | 2.37E-08 |
| Others | | | | 3.08E-04 | 5.59E-05 | 5.52E-07 | 1.23E-02 | 1.87E-12 | 7.46E-01 | 6.60E-32 | 1.45E-01 | 2.06E-02 |
| Streptococcus | | | | | 5.94E-02 | 4.64E-01 | 2.22E-01 | 1.22E-04 | 5.18E-05 | 1.23E-23 | 7.18E-02 | 2.54E-01 |
| Gardnerella vaginalis | | | | | | 4.57E-03 | 9.39E-01 | 1.21E-14 | 3.35E-05 | 9.29E-58 | 4.04E-01 | 7.20E-01 |
| Ca. L. vaginae | | | | | | | 2.27E-01 | 2.09E-10 | 2.23E-07 | 7.58E-52 | 5.13E-02 | 5.38E-01 |
| Atopobium vaginae | | | | | | | | 6.71E-09 | 1.03E-03 | 9.40E-18 | 4.63E-01 | 9.58E-01 |
| Lactobacillus jensenii | | | | | | | | | 1.00E-14 | 2.64E-24 | 7.36E-09 | 1.12E-05 |
| Sneathia amnii | | | | | | | | | | 7.70E-21 | 3.30E-02 | 4.77E-03 |
| No Type | | | | | | | | | | | 5.67E-16 | 1.01E-18 |
| Prevotella bivia | | | | | | | | | | | | 4.67E-01 |
| Mycoplasma | | | | | | | | | | | | |

b

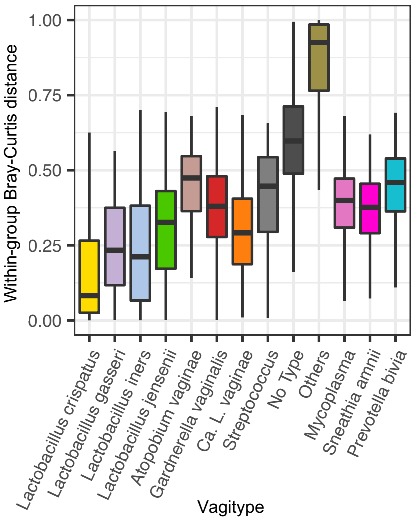


| | Lactobacillus iners | Lactobacillus gasseri | Others | Streptococcus | Gardnerella vaginalis | Ca. L. vaginae | Atopobium vaginae | Lactobacillus jensenii | Sneathia amnii | No Type | Prevotella bivia | Mycoplasma |
|---|---|---|---|---|---|---|---|---|---|---|---|---|
| Lactobacillus crispatus | 0.001 | 0.001 | 0.001 | 0.001 | 0.001 | 0.001 | 0.001 | 0.001 | 0.001 | 0.001 | 0.001 | 0.001 |
| Lactobacillus iners | | 0.001 | 0.001 | 0.001 | 0.001 | 0.001 | 0.001 | 0.001 | 0.001 | 0.001 | 0.001 | 0.001 |
| Lactobacillus gasseri | | | 0.001 | 0.001 | 0.001 | 0.001 | 0.001 | 0.001 | 0.001 | 0.001 | 0.001 | 0.001 |
| Others | | | | 0.001 | 0.001 | 0.001 | 0.001 | 0.001 | 0.001 | 0.001 | 0.001 | 0.001 |
| Streptococcus | | | | | 0.001 | 0.001 | 0.001 | 0.001 | 0.001 | 0.001 | 0.001 | 0.001 |
| Gardnerella vaginalis | | | | | | 0.001 | 0.001 | 0.001 | 0.001 | 0.001 | 0.001 | 0.001 |
| Ca. L. vaginae | | | | | | | 0.001 | 0.001 | 0.001 | 0.001 | 0.001 | 0.001 |
| Atopobium vaginae | | | | | | | | 0.001 | 0.001 | 0.001 | 0.001 | 0.001 |
| Lactobacillus jensenii | | | | | | | | | 0.001 | 0.001 | 0.001 | 0.001 |
| Sneathia amnii | | | | | | | | | | 0.001 | 0.001 | 0.001 |
| No Type | | | | | | | | | | | 0.001 | 0.001 |
| Prevotella bivia | | | | | | | | | | | | 0.001 |
| Mycoplasma | | | | | | | | | | | | |

c

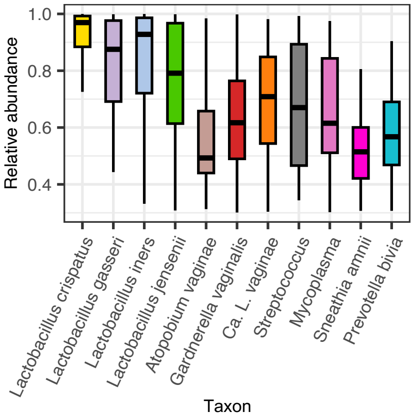


| | Lactobacillus iners | Lactobacillus gasseri | Streptococcus | Gardnerella vagilis | Ca. L. vagie | Atopobium vagie | Lactobacillus jensenii | Sneathia amnii | Prevotella bivia | Mycoplasma |
|---|---|---|---|---|---|---|---|---|---|---|
| Lactobacillus crispatus | 2.95E-10 | 1.74E-05 | 3.73E-13 | 2.03E-103 | 3.23E-63 | 8.69E-17 | 1.51E-06 | 6.99E-26 | 1.87E-15 | 1.81E-13 |
| Lactobacillus iners | | 0.189573097 | 5.11E-08 | 1.10E-85 | 2.39E-40 | 3.06E-13 | 0.011269241 | 5.65E-21 | 2.08E-11 | 1.19E-08 |
| Lactobacillus gasseri | | | 5.00E-05 | 2.93E-17 | 1.58E-09 | 3.18E-10 | 0.15491567 | 1.31E-14 | 1.85E-08 | 1.14E-05 |
| Streptococcus | | | | 0.182155832 | 0.702828676 | 0.009654401 | 0.01397339 | 0.001528026 | 0.085641898 | 0.5279476 |
| Gardnerella vagilis | | | | | 2.27E-06 | 0.006971471 | 3.43E-07 | 0.000211738 | 0.178003246 | 0.5802233 |
| Ca. L. vagie | | | | | | 6.23E-05 | 0.001541758 | 3.93E-08 | 0.001996183 | 0.2666205 |
| Atopobium vagie | | | | | | | 4.63E-07 | 0.975337971 | 0.286088802 | 0.0313794 |
| Lactobacillus jensenii | | | | | | | | 3.64E-09 | 3.20E-05 | 0.0026537 |
| Sneathia amnii | | | | | | | | | 0.168500728 | 0.0086228 |
| Prevotella bivia | | | | | | | | | | 0.1962229 |

**Fig. S1 Shannon index (a), within-group Bray-Curtis distance (b) of the vaginal microbiomes classified by**

**vagitypes and the relative abundance of the predominant taxa in related vagitypes (c).** The *P*-values of the differences pairwise measured by the Wilcoxon test are shown beside the bar plots.

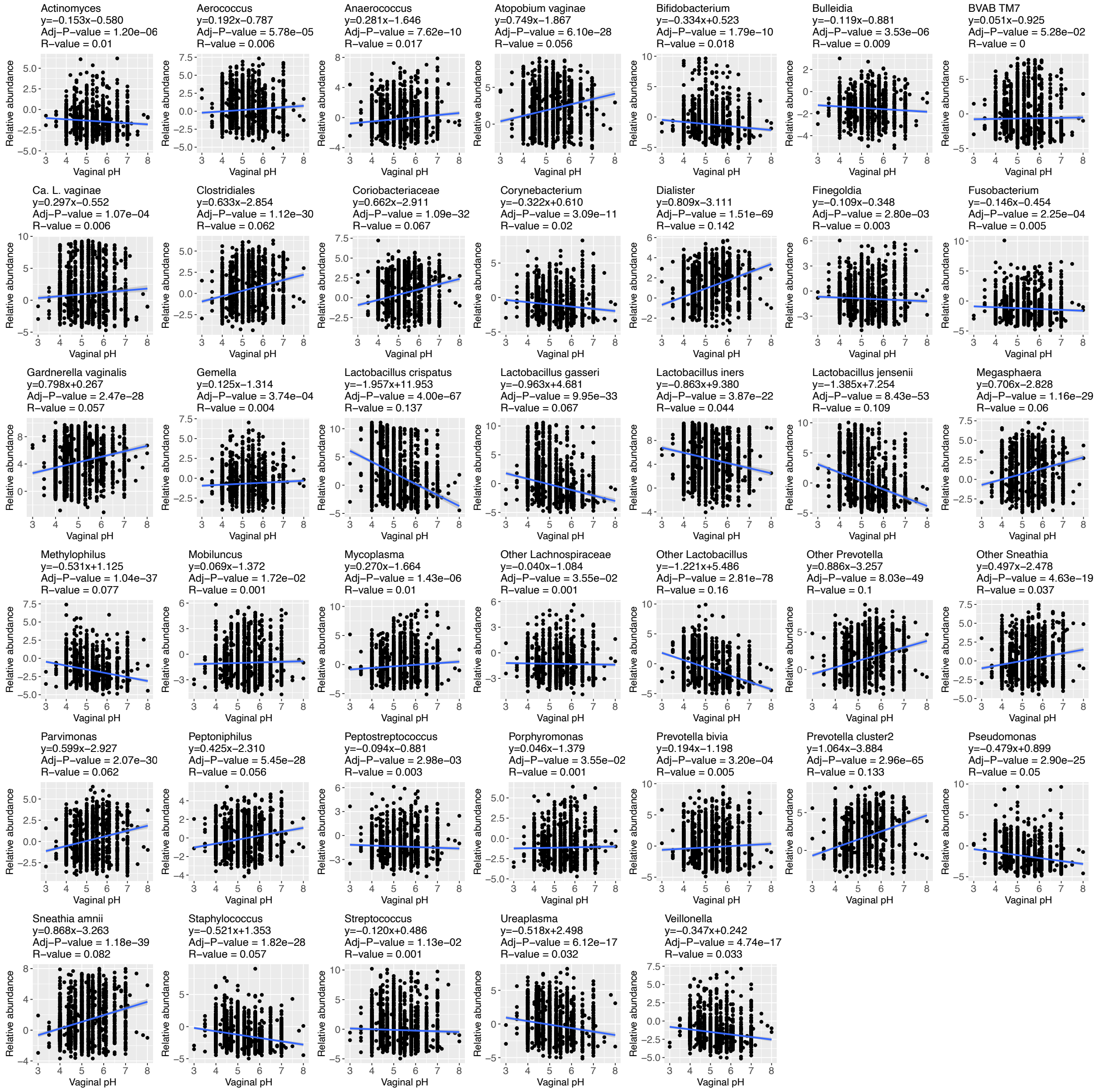


**Fig. S2 Linear relationship between the relative abundance of the vaginal taxa and the vaginal pH tested by the 'lm' function in R.** The *P*-values of the linear correlation were adjusted by the Benjamini-Hochberg Procedure.

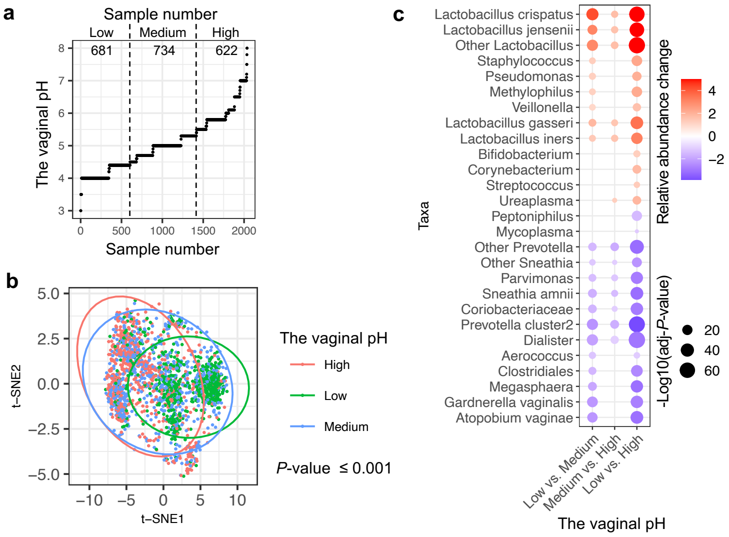


**Fig. S3 The association between the vaginal pH and the vaginal microbiomes. (a)** The classification of the VMBs of the VaHMP project into three groups, i.e., pH low, pH median, and pH high, with similar sample sizes. **(b)** The dissimilarity of the VMBs quantified and visualized by a t-distributed stochastic neighbor embedding plot (t-SNE). The VMBs were color-coded by the vaginal pH levels. The impact of the vaginal pH on the composition of the VMBs was measured by the Adonis test. **(c)** Differential abundance analysis of taxa in the VMBs with different vaginal pH using the 'ADLEx2' package in R. The adjusted *P*-value of relative abundance differences was tested by the 'aldex.ttest' function using the Mann–Whitney U test value, followed by the Benjamini-Hochberg correction. The relative abundance change was measured by the 'aldex.effect' function and quantified by the per-feature median difference between two conditions.

**Table S1 The vaginal pH with a taxon predominant in the VMB.**

| Taxa | The vaginal pH; Median (25th quartile, 75th quartile) | Case number* |
|---|---|---|
| Lactobacillus crispatus | 4.4 (4, 4.7) | 377 |
| Other Lactobacillus | 4.85 (4.7, 5.075) | 4 |
| Bifidobacterium | 4.7 (4.7, 5.375) | 18 |
| Lactobacillus gasseri | 4.7 (4.4, 5) | 51 |
| Lactobacillus iners | 4.7 (4.4, 5) | 606 |
| Pseudomonas | 5 (4.4, 5.75) | 7 |
| Atopobium vaginae | 5 (4.7, 5.8) | 21 |
| Lactobacillus jensenii | 5 (4.7, 5.3) | 34 |
| Other Sneathia | 5.55 (5.1, 5.975) | 4 |
| Fusobacterium | 5.55 (5.075, 6.1) | 4 |
| BVAB TM7 | 5.5 (5.3, 5.8) | 13 |
| Mycoplasma | 5.55 (5.15, 6) | 28 |
| Streptococcus | 5.5 (5, 6.1) | 42 |
| Ca. L. vaginae | 5.3 (5, 5.8) | 228 |
| Gardnerella vaginalis | 5.3 (5, 5.8) | 395 |
| Prevotella cluster2 | 5.8 (5, 5.85) | 20 |
| Other Prevotella | 5.65 (5.3, 5.85) | 20 |
| Prevotella bivia | 5.8 (5, 6.1) | 21 |
| Sneathia amnii | 5.8 (5.3, 5.8) | 38 |
| Anaerococcus | 6.1 (5.3, 6.5) | 5 |
| Gemella | 5.3 (5.3, 5.3) | 1 |
| Veillonella | 5.3 (5.3, 5.3) | 1 |
| Parvimonas | 5.8 (5.8, 5.8) | 1 |
| Porphyromonas | 5.9 (5.9, 5.9) | 1 |
| Finegoldia | 7 (7, 7) | 1 |

| | | |
|---|---|---|
| Ureaplasma | 5.65 (5.575, 5.725) | 2 |
| Corynebacterium | 6.75 (6.625, 6.875) | 2 |
| No Type | 5.5 (5, 5.8) | 92 |

Abbreviation: NA (not applicable)

* Taxa that were predominant in less than three vaginal microbiomes were not classified in the BVT method.

**Movie S1 The third most important component, t-SNE3, was added to the t-SNE plot shown in Fig. 1b. The movie shows the 3D view of the t-SNE plot.**
